# Full-field optical coherence tomography for the diagnosis of giant cell arteritis

**DOI:** 10.1101/2020.05.21.108092

**Authors:** Thomas Maldiney, Hélène Greigert, Laurent Martin, Emilie Benoit, Catherine Creuzot-Garcher, Pierre-Henry Gabrielle, Jean-Marie Chassot, Claude Boccara, Daniel Balvay, Bertrand Tavitian, Olivier Clément, Sylvain Audia, Bernard Bonnotte, Maxime Samson

## Abstract

Histopathological examination of temporal artery biopsy (TAB) remains the gold standard for the diagnosis of giant cell arteritis (GCA) but is associated with essential limitations that emphasize the need for an upgraded pathological process. This study pioneered the use of full-field optical coherence tomography (FF-OCT) for rapid and automated on-site pathological diagnosis of GCA. Sixteen TABs (12 negative and 4 positive for GCA) were selected according to major histopathological criteria of GCA following hematoxylin-eosin-saffron-staining for subsequent acquisition with FF-OCT to compare structural modifications of the artery cell wall and thickness of each tunica. Gabor filtering of FF-OCT images was then used to compute TAB orientation maps and validate a potential automated analysis of TAB sections. FF-OCT allowed both qualitative and quantitative visualization of the main structures of the temporal artery wall, from the internal elastic lamina to the *vasa vasorum* and red blood cells, unveiling a significant correlation with conventional histology. FF-OCT imaging of GCA TABs revealed destruction of the media with distinct remodeling of the whole arterial wall into a denser reticular fibrous neo-intima, which is distinctive of GCA pathogenesis and accessible through automated Gabor filtering. Rapid on-site FF-OCT TAB acquisition makes it possible to identify some characteristic pathological lesions of GCA within a few minutes, paving the way for potential machine intelligence-based or even non-invasive diagnosis of GCA.

## Introduction

Giant cell arteritis (GCA) is a large vessel vasculitis that mainly affects the aorta and the branches of the external carotid, with a predilection for the temporal arteries(1). Even though we now have an accurate understanding of its complex pathogenesis, the causative agent of GCA is still unknown(2). Mostly occurring in northern European females between 50 and 80 years old, the predominant cranial phenotype is usually revealed by new-onset headache, temporal artery tenderness, jaw claudication, and partial or complete visual loss associated with possible systemic symptoms, notably fever, weight loss and weakness(3). The critical complications of GCA include anterior ischemic optic neuropathy, stroke, aortic aneurysm or dissection; these serious complications being responsible for the prognosis of the disease and the need for prolonged high-dose glucocorticoid treatment(4).

The diagnosis of GCA usually relies on the association of concurrent clinical, biological and pathological features of vasculitis that are revealed by temporal artery biopsy (TAB)(5). Significant advances in the field of medical imaging have improved the assessment of the extent of vasculitis and refined non-invasive diagnosis and follow-up(6,7). For instance, the validity of hypoechoic thickening surrounding the temporal artery wall with color duplex sonography (CDS), also referred to as the halo sign, was confirmed at least three times in a meta-analysis for the diagnosis and follow-up of GCA(8). However, the combination of intense infiltration of mononuclear cells in the three layers of the artery, fragmentation of the internal elastic lamina (IEL), intimal hyperplasia and neoangiogenesis on TAB histological examination undoubtedly remains the gold standard for GCA diagnosis in all study group guidelines(9,10).

Apart from rare local complications(11), TAB is a safe procedure(12). Nevertheless, the segmental and focal nature of transmural inflammation in GCA generates skip lesions(13) and is responsible for a significant false-negative rate of up to 30%(14) that makes it necessary to either increase biopsy length(15) or to perform a contralateral TAB(16). These limitations emphasize the potential interest and need for an upgraded pathological procedure dedicated to the diagnosis of GCA.

Based upon optimization of the technology described by Fujimoto and colleagues in the early 1990s,(17,18) full-field optical coherence tomography (FF-OCT) exploits *en face* white-light interference microscopy to provide not only ultra-high resolution images of biological structures(19) but also subcellular metabolic contrast in the tissue depth(20). Until now, most groups have focused on the potential role of FF-OCT during oncologic interventions as new routine approach to surgical pathology(21), and, except for one preliminary study in which the superficial temporal arteries were imaged with dermal OCT(22), there has been no reported attempt to employ high definition interference microscopy for the pathological diagnosis of GCA. The present work pursues the hypothesis that FF-OCT could help both the clinician and pathologist to improve TAB performance, and compares, for the first time, FF-OCT and conventional histological examination for the pathological diagnosis of GCA.

## Materials and Methods

### Preparation of TAB sections

This study was approved by the Institutional Review Board and the Ethics Committee of the Dijon University Hospital. All patients suspected of GCA and scheduled for TAB surgery at the Dijon University Hospital Ophthalmology department from January 2013 to December 2016 were included. All patients provided a signed written informed consent form before inclusion. TAB was performed according to standard procedure, and fresh biopsies were sent to the pathology department. A ten millimeter segment was used for conventional hematoxylin-eosin-saffron (HES) staining, and the other part of the artery segment was immediately frozen at -196°C in fetal bovine serum and dimethyl sulfoxide (10%). Samples were slowly defrosted, the surrounding tissue was removed, and transversal 1 mm-thick sections were cut with a triangular-bladed scalped and placed in complete RPMI culture medium before placement on the sample holder for FF-OCT imaging.

### Histological TAB selection

A total of sixteen TABs were selected for subsequent analysis with optical coherence microscopy. Twelve negative TABs were identified according to the absence of mononuclear cell infiltrate, IEL fragmentation or neoangiogenesis and defined as the control TABs. In these control samples, the pathologist studied the qualitative aspect of the temporal artery wall to distinguish between negative TAB with normal intima (referred to as niTAB.1 to 9, n = 9) and negative TAB with intimal hyperplasia (referred to as ihTAB.1 to 3, n = 3). These control TABs were compared to four specimens that met the major histopathological criteria for the diagnosis of GCA (referred to as gcaTAB.1 to 4, n = 4).

### FF-OCT imaging

FF-OCT images were acquired with a commercially available FF-OCT apparatus (Light-CTScanner, LLTech SAS, Paris, France)(23). Briefly, illumination was provided by a LED source with short coherence length ensuring a sectioning ability or axial resolution of 1 µm. In the FF-OCT set-up, 10x microscope objectives are placed in the interferometer arms in the Linnik configuration, bringing a transverse resolution of 1.5 µm. Following full-field illumination of the axial TAB section, FF-OCT images were captured with a complementary metal oxide semiconductor camera. The theoretical penetration depth for the TAB specimen was approximately 200 µm. The TAB section was placed in the dedicated sample holder with its revolution axis perpendicular to the imaging plane so that one FF-OCT slice showed the architecture of the TAB section from the lumen to the outer wall. A series of FF-OCT slices with 1.5 µm spacing were recorded in depth, and ImageJ 1.52o software was used for axial reconstruction of TAB FF-OCT imaging.

### Image and statistical analysis

Quantitative FF-OCT image analysis and tunica thickness were accessible with a contrast-based ImageJ 1.52o protocol (Plot Profile Function) and calculated as the mean of three representative measurements throughout each TAB section. NDP.view software version 2.6.17, provided by Hamamatsu, allowed similar measurements from scanned glass slides following HES staining. Statistics were calculated using GraphPad Prism version 5. For intima-to media ratios, values reported as medians and interquartile ranges were discriminated by Mann-Whitney tests. The Pearson r coefficient was calculated to evaluate the strength of the linear correlation between histology and FF-OCT measurements of media or intima thicknesses. Interval two-tailed P < 0.05 was considered statistically significant. Finally, orientation maps were calculated for a selection of both healthy and GCA-positive TABs following Gabor filtering of the axial reconstructed images with a custom-made software based on Matlab 2018b (Matworks, Natick, MA).

## Results

### Qualitative FF-OCT imaging

Similar to histological preparation, TAB sections acquired with FF-OCT allow the identification of several important structures within the artery wall (Figure 1). Notably, the tripartite architecture is perceived with a clear separation between intima, media and adventitia (Figure 1A). Interestingly, the physical junction between the intima and media appears as a thin hypo-reflective serpentine band that most obviously corresponds to the IEL (Figure 1A, black arrow). Moreover, in Figures 1B and C obtained with FF-OCT, the *vasa vasorum* can be seen distinctly within the arterial wall and the red blood cells can be identified precisely, returning a spherical contrast highly similar to the one obtained with conventional histology (Figure 1D to F). Indeed, the *vasa vasorum* display similar architecture with both techniques, revealing small (20 to 80 µm) blood vessels characterized by thin elastic walls and a round to oval shape directly inserted into the outlying thread of the temporal artery (*i.e*. mostly between the media and the adventitia layers) (Figures 1B and E, white arrow shows arterial thrombi). Magnification of the lumen of the *vasa vasorum* makes it possible to observe the red blood cells. These cells resemble partially transparent pink-colored ovoid structures following HES staining or a collection of iso-reflective dots with a surrounding hypo-reflective annulus on direct FF-OCT acquisition (Figures 1C and F, black asterisk).

**Figure 1.**
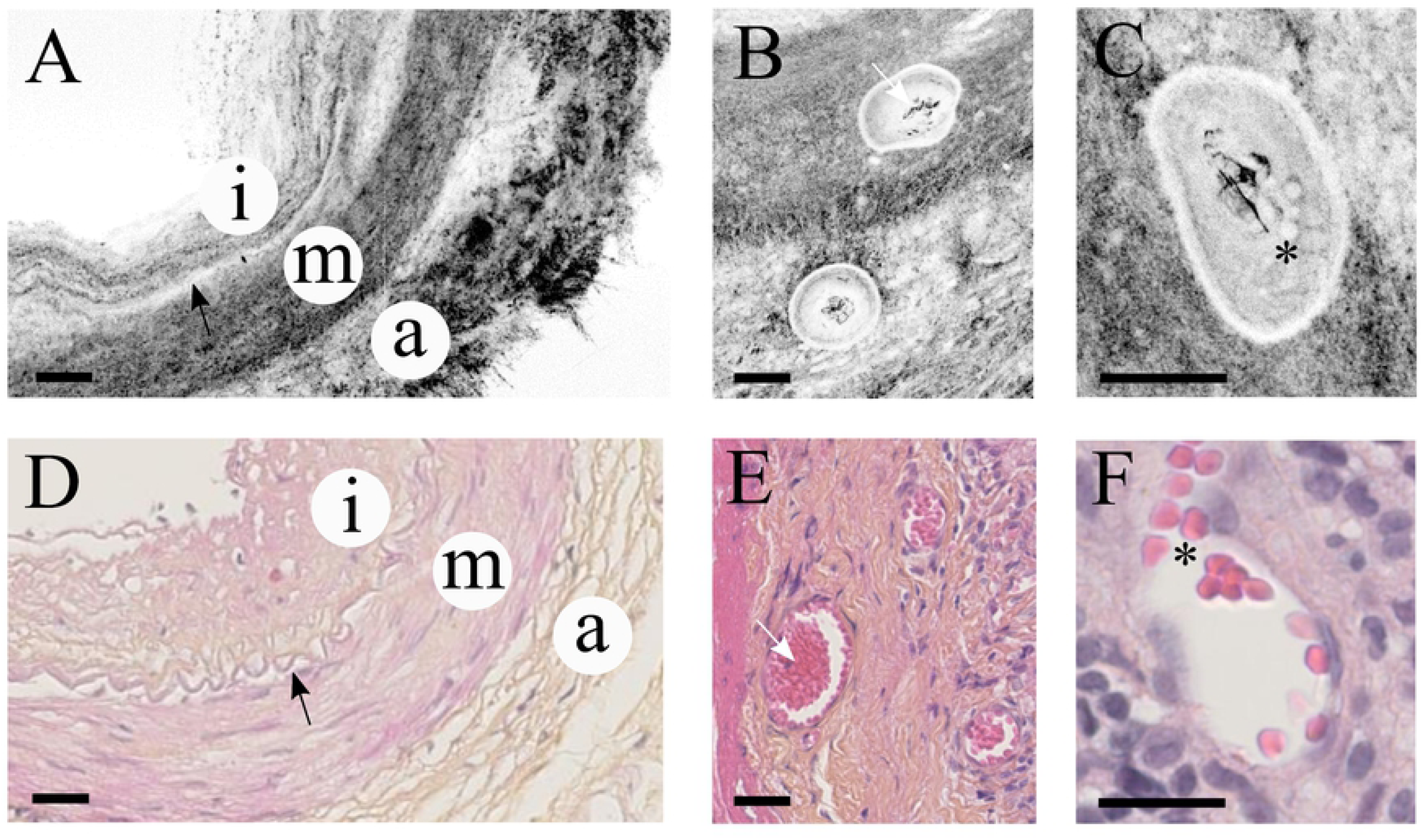
TAB architecture imaging. Arterial wall (A, D), vasa vasorum (B, E) and red blood cells (C, F) imaging with FF-OCT (A to C) and conventional histology following HES staining (D to F). Scale bar represents 50 µm. Black arrow shows IEL. White arrow shows arterial thrombus. Asterisk marks red blood cells. Legend : a, adventitia ; i, intima ; m, media.

FF-OCT acquisition and histological images from negative TAB samples are compared in Figures 2A to D. Figures 2A and B display a representative negative TAB specimen with a thin intimal layer (niTAB). Figures 2C and D show a negative TAB with intimal hyperplasia (hiTAB). No matter the group of negative TAB, the overall architecture of the vessels is preserved, and there is a clear distinction between intima, media and adventitia. Indeed, the tunica media displays a relative hyper-reflectivity on the image acquired with FF-OCT when compared with the tunica intima, whose thin muscle fibers mostly run parallel to the global circular orientation of the TAB section (see magnified region from Figures 2A and C). Similar conclusions regarding the differential contrast and circular symmetry within the two inner layers of the arterial wall separated by the IEL, which appears as a hypo-reflective strip in FF-OCT, can be drawn from the analysis of all negative TAB specimens (Supplemental Figures S1 and S2). In Figures 2A and C we can see that the tunica adventitia is constructed on a denser and more complex fibrous connective tissue that also seems to follow the overall circular symmetry of the system. When compared with Figure 2A, the FF-OCT-acquired TAB section from Figure 2C is characterized by an increased intima thickness (see magnified region). These observations can mostly be transposed for the comparison of all hiTAB specimens detailed in Supplemental Figures S1 and S2. The results obtained with FF-OCT analysis largely correlate with the data obtained after conventional histology (Figures 2B and D). Indeed, the intima in Figure 2B appears much thinner than in Figure 2D in which the intima is thicker than the media.

**Figure 2.**
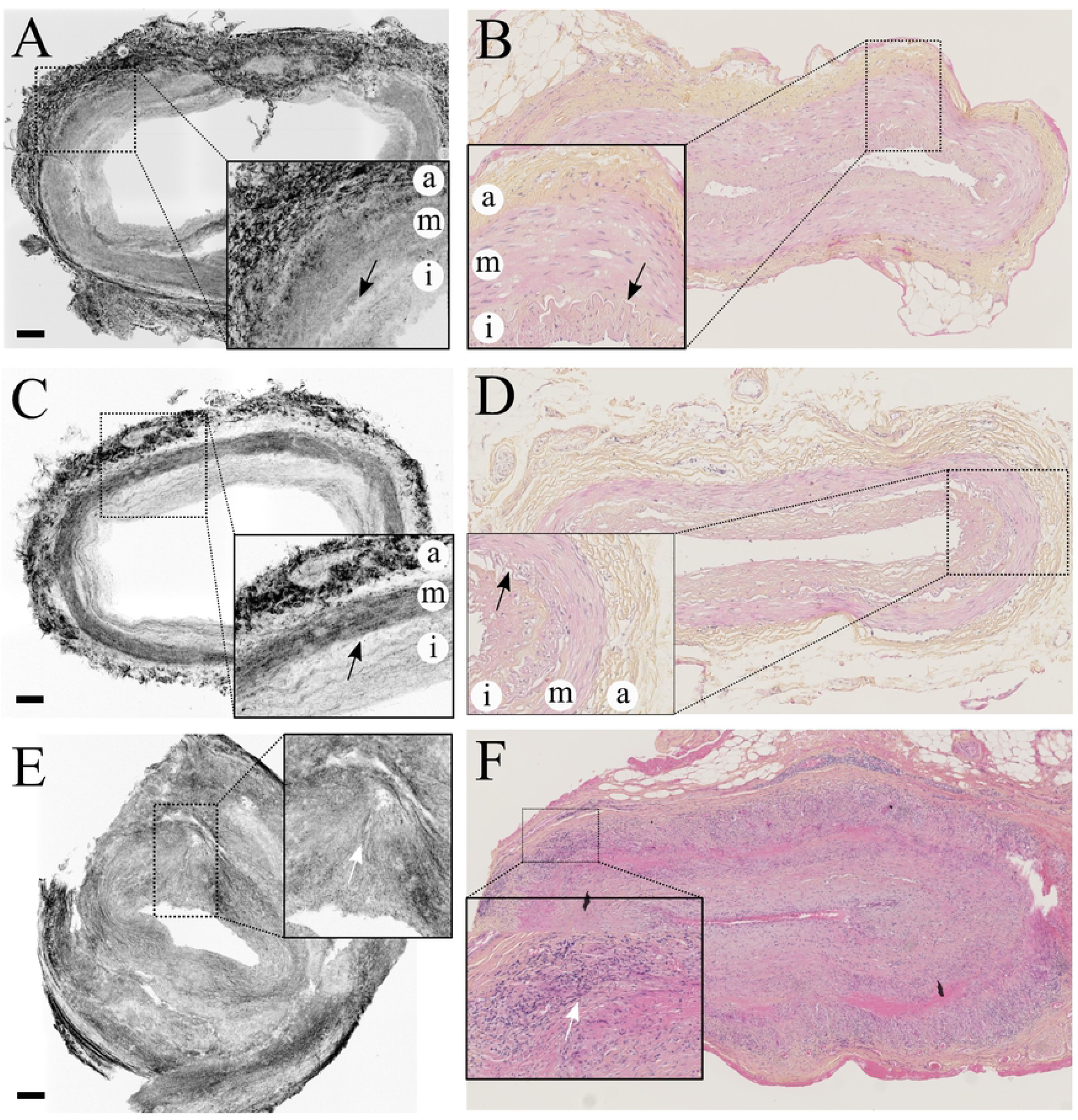
Qualitative imaging of TAB specimens. Comparison of FF-OCT (A, C, E) and conventional histology (B, D, F) imaging. A and B corresponds to niTABs (n = 9), C and D to ihTABs (n = 3), E and F to gcaTABs (n = 4). Scale bar represents 100 µm. Black arrow shows the IEL, white arrow shows rupture of the circular symmetry and mononuclear infiltrate). Legend : a, adventitia ; i, intima ; m, media.

In figures 2E, 2F and S3, the TABs are positive for GCA. The conventional histopathological images show a relatively preserved media that is strongly infiltrated by T-cells, macrophages and multinucleated cells (see the magnified region from Figure 2F). By contrast, FF-OCT acquisition demonstrates a complete disruption of both regular reflectivity and circularity of the media and intima-associated connective tissue fibers due to the infiltration of inflammatory cells. This process remodels the structure of the artery into a denser, reticular, fibrous, collagen-rich structure responsible for both the progressive destruction of the media and the formation of a neo-intima (see the magnified region from Figure 2E). Figure 2E, obtained with FF-OCT, confirms the fragmentation of the internal elastic lamina along with a rebalancing of the contrasts throughout the netlike fibrous structure connecting all three layers. Similar to the corresponding image obtained with conventional histology (Figure 2F), there is no clear distinction between the intima and the media, which is consistent with the stage of the disease. The same conclusion can be drawn from Supplemental Figures S3, which shows the supporting material in which the reticular fibrous neo-intima almost completely obstructs the arterial lumen.

### Quantitative FF-OCT imaging

Given that FF-OCT images provide good quality spatial resolution, we hypothesized that proper image analysis could return quantitative information regarding both the thickness of the artery wall layers and the global architecture of the underlying connective tissue. Figures 3A and B show the main aspects of contrast-based ImageJ protocol along a linear profile drawn across the arterial wall of a negative TAB section (Figure 3A). The protocol was designed to access the most precise measurements for each tunica of the vessel. The gray-scale plot profile from Figure 3B confirms a significant rupture in contrast between the intima and media, as well as between the media and adventitia, allowing concomitant measurements of the thickness of each artery wall layer. Software provided by Hamamatsu facilitated similar measurements from scanned glass slides following HES staining (data not shown). In addition, Gabor filtering was applied to the same reconstructed negative TAB section in order to provide vector orientation maps and subsequent global analysis of the symmetry of the arterial section (Figure 3C). As expected from the previous qualitative analysis, Gabor filtering of FF-OCT-acquired negative TAB section returned a perfect orientation match from one point of the artery to its exact opposite following a 180-degree rotation, as demonstrated by the paired color system respecting an overall 180-degree rotational symmetry. A similar procedure was applied for the analysis of the gray-scale plot profile from positive TAB sections (Figures 3D). Due to a high heterogeneous contrast within the whole TAB section of GCA patients, Figure 3E shows almost no possible distinction between the different layers composing the artery wall with a contrast oscillating between 500 and 2000 arbitrary units from the very inner to the outer layer. Subsequent Gabor filtering of the positive section proves the pathological loss of the 180 degree rotational symmetry-based vector orientation match, as illustrated by the relative chaotic color distribution within the layers of the artery resulting in a rainbow-like appearance (Figure 3F).

**Figure 3.**
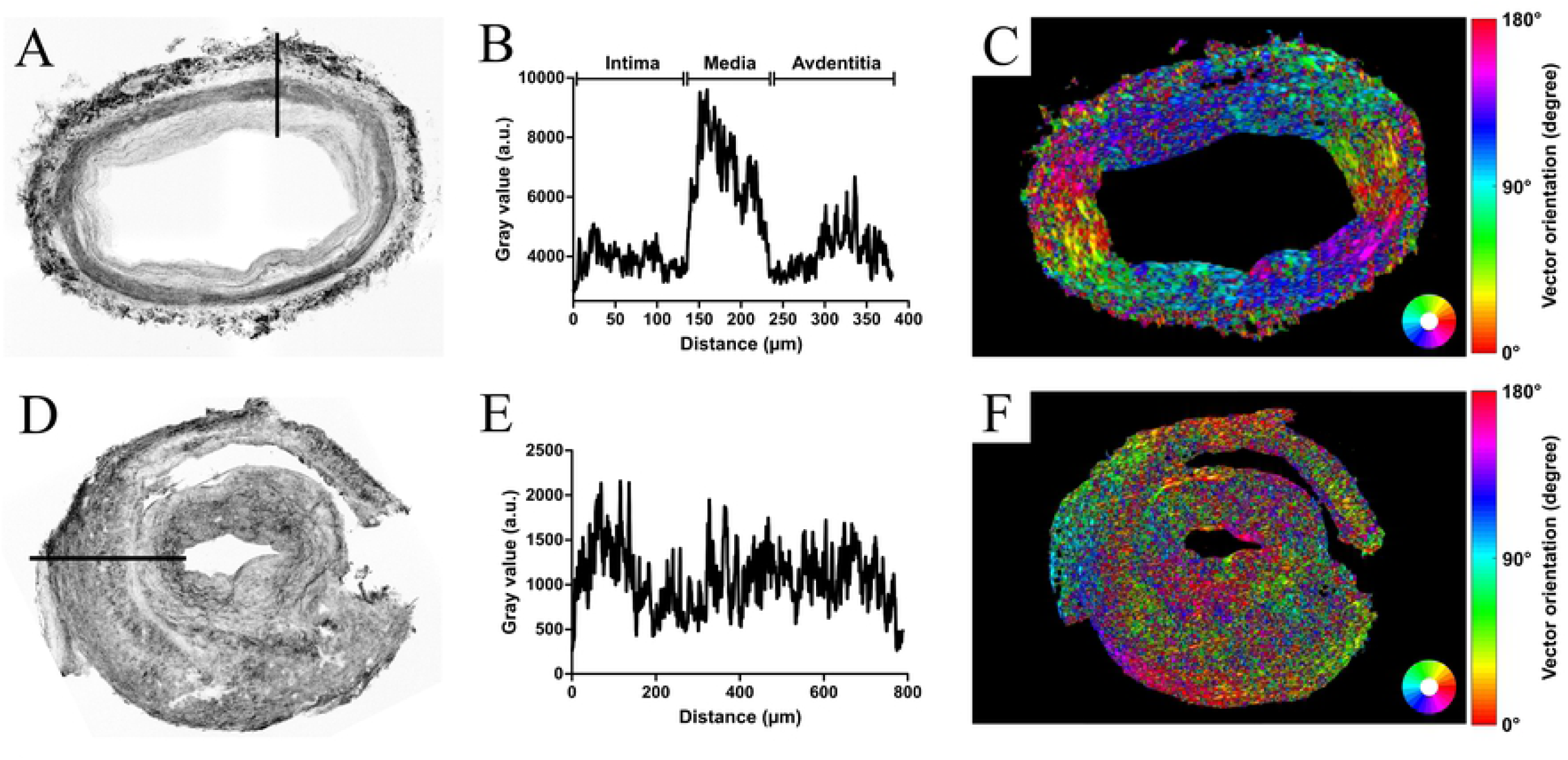
Image analysis of TAB sections. ihTAB1 FF-OCT translational section for ImageJ graphical plot (A). ImageJ graphical plot of ihTAB1 FF-OCT translational section (B). Orientation maps after Gabor filtering of ihTAB1 FF-OCT translational section (C). gcaTAB3 FF-OCT translational section for ImageJ graphical plot (D). ImageJ graphical plot of gcaTAB3 FF-OCT translational section (E). Orientation maps after Gabor filtering of gcaTAB3 FF-OCT translational section (F).

Figure 4A shows an example of a negative TAB for which FF-OCT images and conventional histology match perfectly from the inner to the outer layer of the biopsy. Regardless of intima thickness or GCA status (when quantifiable for GCA patients), quantitative analysis of both intima and media thickness confirms the absence of statistical difference between FF-OCT-based and histology-based measurements of intima-to-media ratios (Figure 4B). Moreover, these results let to an accurate association of both quantitative classification and qualitative selection established by the pathologist for negative sections. Indeed, TABs with thin intima (Supplemental Figure S1) and TABs with intimal hyperplasia (Supplemental Figure S2) consequently appear as two separate groups: one with normal intima/media ratio (I/M) <1 and another that shows an intimal hyperplastic response with I/M between 1 and 2. When data were accessible for TABs with GCA lesions, quantification brought out a third GCA group defined by I/M largely > 2. Finally, data from Figures 4C and D demonstrate a significant correlation between the thickness of the intima (Figure 4C) and media (Figure 4D) measured with FF-OCT and conventional histology.

**Figure 4.**
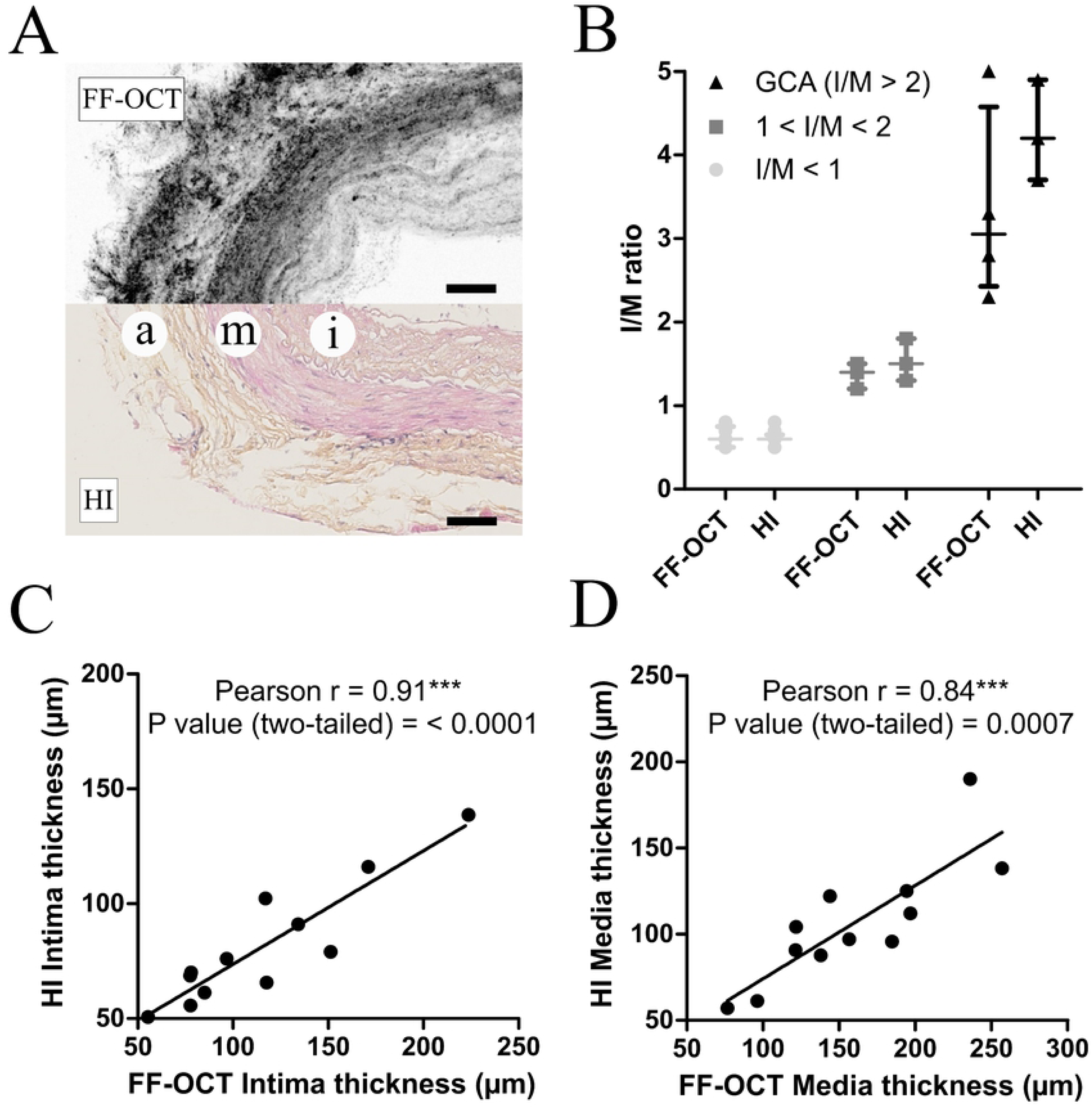
Quantitative analysis of TAB sections. FF-OCT and conventional histology cross-sectional match for the visualization of temporal artery wall (A). Comparison of FF-OCT and conventional histology-based intima-to-media ratios for both healthy and GCA-positive TAB sections (B). Correlation curves between FF-OCT and conventional histology for the measurement of intima (C) and media (D) thickness. Scale bar represents 100 µm.

## Discussion

The present work describes the first attempt to assess the potential of FF-OCT for the diagnosis of GCA in comparison with conventional histology. The first advantage of FF-OCT is that it provides rapid (within minutes) and on-site acquisition of TAB sections. We demonstrate, from the analysis of both healthy and GCA-positive TAB sections, that the high spatial resolution of FF-OCT technology makes it possible to visualize with precision several essential structures correlated with the diagnosis of GCA. Notably, we found that FF-OCT accurately returns both qualitative and quantitative information relative to the structure of the three arterial tissue layers and the internal elastic lamina or *vasa vasorum*, with a significant correlation to histopathological imaging. When focusing on the FF-OCT analysis of healthy TAB sections, the inverted intima-to-media ratio can be interpreted as a reflection of the stage in human atherosclerosis(24). Moreover, we provide preliminary proof that automated Gabor filtering could deliver both immediate and essential structural information regarding the preservation of the regular circularity of the media and intima-associated connective tissues, paving the way for potential machine intelligence-based pathological diagnosis of GCA. FF-OCT acquisitions return additional and complementary information with focus on the appearance and structural orientation of the underlying fibrous supporting tissue within each layer of the temporal artery. When TABs from GCA patients were compared to the global circular symmetry of healthy TAB sections, FF-OCT imaging revealed the destruction of the media layer and the modification of the arterial wall structure, which was rearranged into a denser reticular fibrous neo-intima, distinctive of GCA pathogenesis(25). Despite the current success of non-invasive techniques like CDS, a precise FF-OCT-based analysis of the temporal artery wall on a meso-structural level remains of particular importance for the diagnosis of GCA. There is, however, a potential pitfall for GCA diagnosis with CDS since the atherosclerotic lesions responsible for significant increases in the thickness of the intima might mimic the halo sign, resulting in false positives(26).

We acknowledge several limitations that had an impact on the use of FF-OCT for rapid on-site pathological diagnosis of GCA in the current study. First, T-cells, macrophages and multinucleated cells, which are hallmarks of GCA, were not visible in the present set-up of FF-OCT, which used previously frozen TAB samples. Despite the high spatial resolution, the loss of information was due to the structural nature of contrast imaging, rendering direct black and white photographs of the specimen without any preparation or staining. However, this limitation is for the most part the result of using defrosted TAB samples with dead cellular material. This issue can be overcome by performing dynamic FF-OCT acquisition of fresh TAB sections, yielding complementary subcellular contrast(27) and putative direct visualization of inflammatory infiltrates. The potential of *en face* white-light interference microscopy demonstrated in this work should encourage further investigations into the FF-OCT-based handheld acquisition probe(28), a promising technology dedicated to direct transcutaneous imaging and further non-invasive diagnosis of GCA.

In conclusion, this preliminary study is the first to compare FF-OCT imaging to the gold standard histopathological procedure for the diagnosis of GCA. It brings conclusive proof regarding the potential of FF-OCT for both qualitative and quantitative structural visualization of underlying fibrous tissues and architectural changes in the arterial wall that occur throughout the inflammatory processes of GCA. After this first promising step, further investigations are warranted to confirm the potential of FF-OCT technology for rapid, on-site, non-invasive diagnosis of GCA.

## Acknowledgments

The authors thank Suzanne Rankin for her help in proofreading the article ; Marion Ciudad, Marine Thébault, Claudie Cladière, Claire Gérard, Mathilde Charlot, Claire Boillin, Martine Breton, Amandine Esnoult, Thibault Ghesquière and Laetitia Barbier for their help in collecting samples ; Céline Shaeffer, Eva Michaud and the Centre de Ressources Biologiques ; Ferdinand Cabanne for their help in the conservation of frozen samples.

## Statement of author contributions

T. M. designed research, performed research, analyzed data and wrote the paper. H. G. performed research. L. M. analyzed data. E. B. performed research and analyzed data. C. C.-G. analyzed data. PH. G. analyzed data. JM. C. performed research and analyzed data.

C. B. designed research and performed research. D. B. performed research and analyzed data. B. T. analyzed data. O. C. analyzed data. S. A. designed research. B. B. designed research, analyzed data, and wrote the paper. M. S. designed research, performed research, analyzed data and wrote the paper.

M. S. is the guarantor of this work and, as such, had full access to all of the data in the study and takes responsibility for the integrity of the data and the accuracy of the data analysis.

## Supplementary Figure captions

**Figure S1. Qualitative imaging of healthy TAB specimens with thin intima layer** (n = 9). Comparison of FF-OCT (A, C, E, G, I, K, M, O, Q) and conventional histology (B, D, F, H, J, L, N, P, R) imaging. A and B correspond to niTAB1, C and D to niTAB2, E and F to niTAB3, G and H to niTAB4, I and J to niTAB5, Kand L to niTAB5, M and N to niTAB7, O and P to niTAB8, Q and R to niTAB9.

**Figure S2. Qualitative imaging of healthy TAB specimens with significant intimal hyperplasia** (n = 3). Comparison of FF-OCT (A, C, E) and conventional histology (B, D, F) imaging. A and B correspond to the ihTAB1, C and D to ihTAB2, E and F to ihTAB3.

**Figure S3. Qualitative imaging of GCA TAB specimens** (n = 4). Comparison of FF-OCT (A, C, E, G) and conventional histology (B, D, F, H) imaging. A and B correspond to gcaTAB1, C and D to gcaTAB2, E and F to gcaTAB3, G and H to gcaTAB4.

